# SurfNet: Reconstruction of Cortical Surfaces via Coupled Diffeomorphic Deformations

**DOI:** 10.1101/2025.01.30.635814

**Authors:** Hao Zheng, Hongming Li, Yong Fan

**Affiliations:** Center for Biomedical Image Computing and Analytics, Philadelphia, PA 19104, USA; Department of Radiology, University of Pennsylvania, Philadelphia, PA 19104, USA; School of Computing and Informatics, University of Louisiana at Lafayette, Lafayette, LA 70503, USA

**Keywords:** Cortical surface reconstruction, diffeomorphic deformation, ODE and thickness estimation

## Abstract

To achieve fast and accurate cortical surface reconstruction from brain magnetic resonance images (MRIs), we develop a method to jointly reconstruct the inner (white-gray matter interface), outer (pial), and midthickness surfaces, regularized by their interdependence. Rather than reconstructing these surfaces separately without taking into consideration their interdependence as in most existing methods, our method learns three diffeomorphic deformations jointly to optimize the midthickness surface to lie halfway between the inner and outer cortical surfaces and simultaneously deforms it inward and outward towards the inner and outer cortical surfaces, respectively. The surfaces are encouraged to have a spherical topology by regularization terms for non-negativeness of the cortical thickness and symmetric cycle-consistency of the coupled surface deformations. The coupled reconstruction of cortical surfaces also facilitates an accurate estimation of the cortical thickness based on the diffeomorphic deformation trajectory of each vertex on the surfaces. Validation experiments have demonstrated that our method achieves state-of-the-art cortical surface reconstruction performance in terms of accuracy and surface topological correctness on large-scale MRI datasets, including ADNI, HCP, and OASIS.^1^

## 1 Introduction

Cortical surface reconstruction (CSR) from structural MRI is widely employed in imaging studies of neurodegenerative diseases (Bertoux et al., 2019; Crutch et al., 2012; Roe et al., 2021) and psychological disorders (Rimol et al., 2012) for quantitatively characterizing cortical morphology. Anatomically, the human cerebral cortex has the topology of a 2D convoluted sheet (i.e., a closed manifold without handles or holes) (Fischl, 2012), bounded by an inner (white matter) surface between the cortical gray matter (GM) and white matter (WM) of the brain and an outer (pial) surface between the cerebrospinal fluid (CSF) and the GM. Due to the partial volume effect in brain MRIs and the highly-folded patterns of the cerebral cortex, voxel-based segmentation methods cannot accurately capture its complex morphology (Bongratz et al., 2022; Fischl, 2012). Well-established CSR methods, such as FreeSurfer (Fischl, 2012), BrainSuite (Shattuck & Leahy, 2002), and the HCP structural pipeline (Glasser et al., 2013), successfully reconstruct cortical surfaces represented as triangular meshes from brain MRIs. However, these methods suffer from high computational costs and lengthy processing times because of their reliance on multiple computationally intensive geometric or image processing algorithms. Moreover, these methods may require manual editing to achieve subvoxel accuracy. For instance, FreeSurfer takes an average of 6 hours to reconstruct the cortical surfaces for a single subject (Fischl, 2012).

Recent deep learning (DL) based CSR methods (Bongratz et al., 2022; Cruz et al., 2021; Hong et al., 2021; Hoopes et al., 2022; Lebrat et al., 2021; Q. Ma et al., 2021, 2022; Ren et al., 2022; Santa Cruz et al., 2022; Zheng et al., 2023b) train neural networks in an end-to-end manner to predict cortical surfaces and have achieved high accuracy while being *hundreds to thousands of times faster* than conventional methods in inference. These methods can be categorized according to the representation of output surfaces: *(I) Implicit surface representation*: The signed distance function (Cruz et al., 2021; Hong et al., 2021), occupancy field (Cruz et al., 2021), and level sets (Ren et al., 2022) are first predicted by neural networks; and the cortical surface is then generated via the Marching Cubes algorithm (Lewiner et al., 2003). *(II) Explicit surface representation*: These methods take as input a coarse initial mesh (e.g., a sphere (X. Chen et al., 2023), over-smoothed convex hull, or group template (Bongratz et al., 2022; Lebrat et al., 2021; Santa Cruz et al., 2022)) or a fine initial mesh (e.g., an existing WM surface (Hoopes et al., 2022; Q. Ma et al., 2021) or one extracted from segmentation maps (Q. Ma et al., 2022; Zheng et al., 2023b)) and directly predict a target mesh.

However, existing *DL-based CSR* methods have several limitations. *First*, the *interdependence* between the inner and outer cortical surfaces is generally ignored. Separate or multistage DL models (Q. Ma et al., 2022; Santa Cruz et al., 2022) are typically trained to reconstruct the inner and outer cortical surfaces, and intersections between them may occur. Simultaneous reconstruction methods (Bongratz et al., 2022; Zheng et al., 2023b) lack topological constraints, resulting in loosely coupled surfaces. *Second*, using a coarse mesh template (e.g., sphere, convex-hull, or over-smoothed template (Bongratz et al., 2022; X. Chen et al., 2023; Dale et al., 1999; Santa Cruz et al., 2022)) as an initialization surface incurs the difficulty of learning large deformations for highly folded cortical regions and may lead to non-smooth deformation and undesirable artifacts. *Third*, no existing DL-based CSR method simultaneously estimates, necessitating a *separate* step to compute cortical attributes as in conventional pipelines (Fischl, 2012; Wu et al., 2021). Incorporating this anatomical constraint could strengthen the inter-surface relationships, enable direct cortical thickness analysis, and enforce topological correctness in the reconstructed surfaces. *Fourth*, existing methods rely on numerical solutions that either use explicit integration over voxel-wise features extracted from convolutional neural networks (CNNs) (Lebrat et al., 2021) or implicit neural ODEs (NODEs) (R. T. Chen et al., 2018) learned from local image subvolumes (Q. Ma et al., 2022). At present, a unified framework seamlessly integrating both approaches does not exist.

We propose a DL-based CSR method, referred to as SurfNet, to *simultaneously* reconstruct both the WM and pial surfaces while directly estimating the cortical thickness by optimizing and deforming an initialization midthickness surface, with ordinary differential equations (ODEs) being leveraged to model the diffeomorphic deformations to ensure topological correctness. First, a fine midthickness surface mesh of genus 0 that lies approximately halfway between the WM and pial surfaces is generated from a cortical ribbon segmentation result and refined by a fast topology correction procedure. Then SurfNet simultaneously learns three diffeomorphic deformations from a multi-channel input, consisting of a 3D MRI image and an initialization surface embedded in a 3D map as a signed distance function, to deform the initialization surface towards the midthickness surface and simultaneously deform it towards inner and outer cortical surfaces, respectively. Specifically, we showcase two ways to model the diffeomorphic deformations: 1) by computing the deformation from a dense stationary velocity field prediction, and 2) by parameterizing the continuous dynamics of the vertex trajectory using neural ODEs. Our framework is flexible to accommodate both the solutions and multiple cortical surfaces are *coupled* in the network optimization. To further enhance the spherical topology of reconstructed surfaces, we propose two regularization terms based on the non-negativeness of the cortical thickness and the symmetric cycle-consistency of the midthickness surface’s deformations. Moreover, a vertex-wise thickness estimation can be obtained by tracing the geodesic trajectory of each vertex during the mesh deformation process.

This paper extends our preliminary work (Zheng et al., 2023a) by proposing a framework that accommodates both volume-wise and vertex-wise velocity fields for learning diffeomorphic deformations, while also expanding the analyses to include more datasets with multiple settings. Specifically, we provide a comprehensive analysis of the CSR framework in three key aspects: 1) *how to initialize* the initialization surface; 2) *how to deform* the initialization surface to jointly reconstruct *multiple* cortical surfaces; and *3)* deformation computation. Extensive evaluations are performed on multiple datasets with detailed ablation studies. In summary, our contributions are threefold:

- We propose SurfNet, a united framework for *coupled* cortical surface reconstruction based on *diffeomorphic* deformation, with simultaneous *cortical thickness estimation*.
- We introduce *two* approaches to model diffeomorphic deformations, showcasing the flexibility of SurfNet.
- We perform extensive experiments on three large public datasets (ADNI (Jack Jr et al., 2008), HCP (Van Essen et al., 2013), and OASIS (Marcus et al., 2007)) and demonstrate SurfNet’s superior performance compared to state-of-the-art alternatives (Bongratz et al., 2022; Cruz et al., 2021; Dong et al., 2024; Lebrat et al., 2021; Q. Ma et al., 2021, 2022; Santa Cruz et al., 2022; Zheng et al., 2023b).

## 2 Related Works

### 2.1 Learning-based Surface Reconstruction

Recent progress in deep geometric learning (Fahim et al., 2021; Gupta & Chandraker, 2020; B. Ma et al., 2021; Park et al., 2019; Sinha et al., 2017; N. Wang et al., 2020; C. Wen et al., 2023; X. Wen et al., 2022; Xie et al., 2020) has facilitated applications in the field of biomedical object reconstruction (Kong & Shadden, 2022; Kong et al., 2021; Pak, Liu, Ahn, et al., 2021; Pak, Liu, Kim, et al., 2021; Sun et al., 2022; X. Wang et al., 2023; Wickramasinghe et al., 2020; Ye et al., 2023), such as aortic valves, liver, hippocampus, pancreas, lung, and whole heart. They can be categorized into two groups. One group utilizes CNNs to learn a deformation field to deform an initialization mesh to the target surface, such as MTM (Pak, Liu, Ahn, et al., 2021). The other group adopts graph neural networks (GNNs)/fully-connected networks (FC-Nets), some combined with CNNs, to deform a coarse mesh template (e.g., ellipsoid, smoothed convex hull) to the target surface. Particularly, Voxel2Mesh uses an adaptive mesh unpooling method (Wickramasinghe et al., 2020), MeshDeformNet combines multiple anatomical structures (Kong et al., 2021), NDF models shapes as conditional deformations of a template deep implicit function (Sun et al., 2022), HeartDeformNets adopts a series of GNN-based deformation blocks (Kong & Shadden, 2022), and NDM integrates parameter function-based primitives and neural diffeomorphic flow parameterized local deformation (Ye et al., 2023). Nevertheless, the cerebral cortex has a more complex shape, and it merits further investigation on how to initialize the template surface and preserve the diffeomorphism of the surface deformation.

The existing DL-based CSR methods try to find a better initialization surface (Q. Ma et al., 2022; Santa Cruz et al., 2022; Zheng et al., 2023b) and deform the initial surface to a target surface by learning a volumetric velocity field (X. Chen et al., 2023; Hoopes et al., 2022; Lebrat et al., 2021; Santa Cruz et al., 2022; Zheng et al., 2023b) or vertex-wise displacement (Bongratz et al., 2022; Hoopes et al., 2022; Q. Ma et al., 2021, 2022). A detailed comparison of these methods is summarized in Table 1. One group of methods represents the surface *implicitly* as a signed distance function (Cruz et al., 2021; Hong et al., 2021), occupancy field (Cruz et al., 2021), or level set (Ren et al., 2022), followed by the marching cubes algorithm (Lewiner et al., 2003) to extract the zero level-set to reconstruct the cortical surfaces. These methods typically rely on computationally intensive topology correction to guarantee the spherical topology of the reconstructed surfaces, and CSR accuracy is also constrained by the volumetric representations. The other group of methods directly deforms an initial mesh (coarse or fine *explicit* mesh templates) to a target mesh (Bongratz et al., 2022; Hoopes et al., 2022; Lebrat et al., 2021; Q. Ma et al., 2021, 2022; Santa Cruz et al., 2022; Zheng et al., 2023b), and the desired mesh topology is maintained by enforcing the deformation to be diffeomorphic through the utilization of regularization terms. For example, PialNN (Q. Ma et al., 2021) deforms the fine WM surface to the pial surface by learning the displacement for each vertex from locally sampled cubes with an FC-Net, followed by Laplacian mesh smoothing. TopoFit (Hoopes et al., 2022) utilizes a series of GNN blocks to predict deformations that warp a coarse template mesh to fit the fine target mesh with upsampling between each deformation step and employs a hinge-spring term to maximize the angle between neighboring triangles. Vox2cortex (Bongratz et al., 2022) incorporates an inter-mesh feature exchange mechanism and a curvature-weighted loss function in a GNN-based deformation network and adopts mesh smoothness and edge length regularizations. Besides relying on regularization terms alone to reduce the mesh self-intersections, other works learn the continuous diffeomorphic flow and model the mapping between vertices as ODEs, as seen in methods like CorticalFlow (Lebrat et al., 2021) and CorticalFlow++ (Santa Cruz et al., 2022), or employ NODEs (R. T. Chen et al., 2018), as demonstrated in CortexODE (Q. Ma et al., 2022). However, these methods neglect the interdependence between the inner and outer cortical surfaces. As a result, they require training multiple models to reconstruct each cortical surface separately and lack essential constraints to ensure non-negative cortical thickness, increasing the risk of generating intersecting surfaces. In contrast, our approach distinguishes itself from existing DL-based CSR methods by enabling the coupled reconstruction of multiple surfaces with explicit thickness constraints.

**Table 1:** A comparison of existing DL-based CSR methods and ours. For input, Vol.: whole 3D MRI volume; cube: sub-volume cropped from the MRI volume; FS: fine surface mesh with detailed structures; CS: coarse surface mesh (e.g., convex-hull or oversmoothed group template); IS: implicit surface representation. For output, explicit surface: triangular mesh; implicit surface: signed distance function or occupancy field. Thickness refers to *direct* cortical thickness estimation. Diffeomorphism refers to diffeomorphic deformation in CSR.

| Methods | Output surf. | Input Img. | Surf. | Coupled CSR | Thickness | Diffeomorphism |
| --- | --- | --- | --- | --- | --- | --- |
| DeepCSR (Cruz et al., 2021)<br>vox2surf (Hong et al., 2021)<br>FastCSR (Ren et al., 2022) | Implicit | vol.<br>vol.+cube<br>vol. |  |  |  |  |
| PialNNQ. Ma et al., 2021 | Explicit | cube | FS |  |  |  |
| CF (Lebrat et al., 2021; Santa Cruz et al., 2022) |  | vol. | CS |  |  | ✓ |
| SurfFlow (X. Chen et al., 2023) |  | vol. | CS |  |  | ✓ |
| vox2cortex (Bongratz et al., 2022) |  | vol. | CS | ✓ |  |  |
| TopoFit (Hoopes et al., 2022) |  | vol. | CS |  |  |  |
| CortexODEQ. Ma et al., 2022 |  | cube | FS |  |  | ✓ |
| SurfNN (Zheng et al., 2023b) |  | vol. | FS+IS | ✓ | ✓ |  |
| ccfCSR (Dong et al., 2024) |  | vol. | FS+IS | ✓ |  | ✓ |
| SurfNet (Ours) |  | vol. | FS+IS | ✓ | ✓ | ✓ |

### 2.2 Diffeomorphic Deformation

Diffeomorphic deformation is a smooth and invertible spatial transformation (Ruelle & Sullivan, 1975). It has been widely used in the modeling and analysis of brain morphometry. LDDMM (Beg et al., 2005) computes a time-dependent velocity vector field based on an ODE. A stationary velocity field (SVF) in conjunction with the scaling and squaring method has been developed to reduce computational complexity (Arsigny et al., 2006). Learning-based methods (Balakrishnan et al., 2019; Li et al., 2022; Mok & Chung, 2020) have been developed to improve computational efficiency. Regularizations such as smoothness (Balakrishnan et al., 2019) and inverse-consistency (Mok & Chung, 2020), have also been introduced to enhance the diffeomorphic property of the deformation. The diffeomorphic deformation strategy has also been adopted in CSR methods. Particularly, CoticalFlow methods (Lebrat et al., 2021; Santa Cruz et al., 2022) propose to solve the ODE vertex-wisely and derive a numerical condition to ensure the homeomorphism of integration by training a *chain* of diffeomorphic deformation models in *sequential* stages to deform an initialization mesh template of a convex hull to the target surface in a coarse-to-fine manner. Recently, NODEs (R. T. Chen et al., 2018) have been adopted to model diffeomorphic flows (Gupta & Chandraker, 2020; Q. Ma et al., 2022; Sun et al., 2022). In particular, CortexODE (Q. Ma et al., 2022) parameterizes the trajectories of vertices on the surface as ODEs and proposes a pipeline to initialize and reconstruct the WM and pial surfaces *sequentially*. The sufficient condition for diffeomorphism can be satisfied if a deformation network is Lipschitz continuous and a sufficiently small step size is used in numerical approximation. In this study, we introduce topology-preserving and inverse-consistent transformation regularizations to enforce diffeomorphic deformation, and demonstrate that our framework accommodates the *two* primary diffeomorphic deformation paradigms: CNN-based and NODE-based approaches.

### 2.3 Cortical Thickness Estimation

There are two types of methods for estimating the cortical thickness (CTh). One is based on the reconstructed WM and pial surfaces. For example, FreeSurfer averages the distance from a point on the WM surface to the closest point on the pial surface and the distance from that point back to the nearest point on the WM surface (Dale et al., 1999; Fischl & Dale, 2000). The other is based on a dense deformation of ribbon segmentation maps. Specifically, DiReCT (Das et al., 2009) estimates the CTh from a spatial deformation velocity field that registers WM segmentation map and a combined WM+GM segmentation map. CortexMorph (McKinley & Rummel, 2023) utilizes a volume registration network to achieve fast calculation of diffeomorphisms for DiReCT on the GPU (4.3s±0.71s per subject). The former requires an additional step to build point correspondence between a pair of reconstructed cortical surfaces, while the latter necessitates training a separate model for dense deformations. In this work, we integrate the CTh estimation into the CSR framework by introducing a deformation trajectory loss, which is seamlessly computed alongside CSR with negligible additional cost and enhances surface accuracy through regularization.

## 3 Methodology

SurfNet, the coupled CSR framework as illustrated in Fig. 1, is presented in the following subsections: coupled learning of cortical surfaces (3.1), loss functions (3.2), modeling of diffeomorphic deformation (3.3), and implementations (3.4).

**Figure 1:**
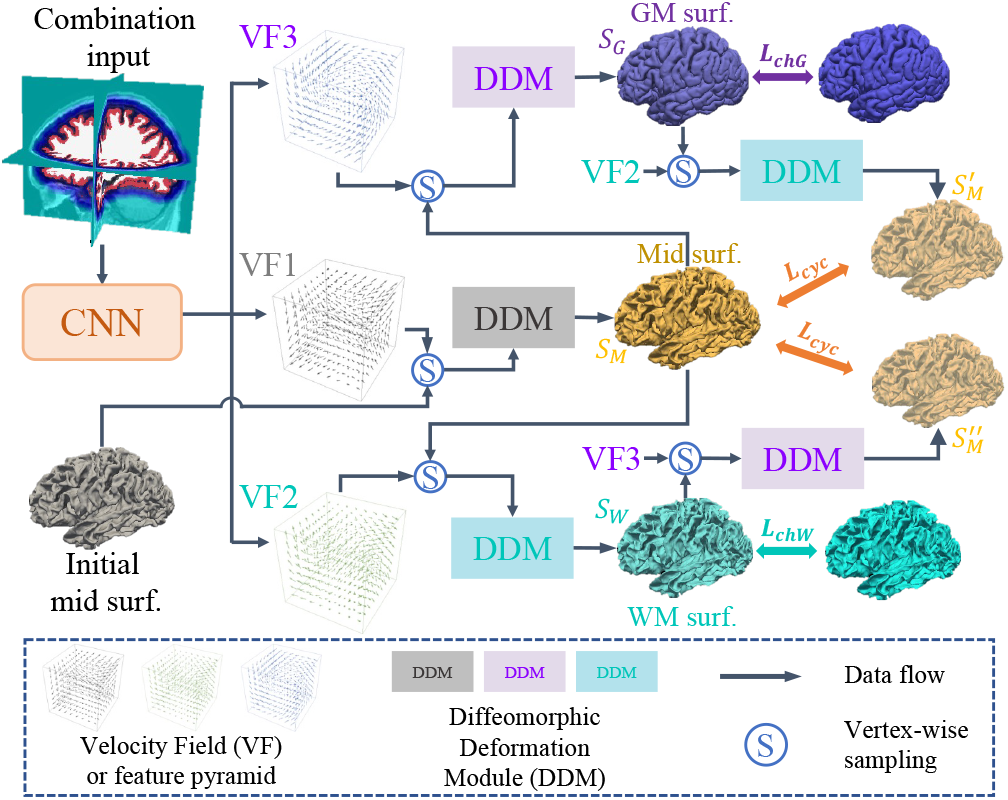
SurfNet for coupled cortical surface reconstruction. Taking as input a combination of MRI brain image, cortical ribbon segmentation maps, and a signed distance map of the midthickness surface, SurfNet learns three diffeomorphic deformations simultaneously to optimize the initial midthickness surface 𝒮_0_ to align with the target midthickness surface 𝒮_*Mid*_ with a diffeomorphic deformation model (DDM), and deform 𝒮 _*Mid*_ outward and inward towards the pial surface 𝒮 _*G*_ and the WM surface 𝒮 _*W*_ with another two DDMs, respectively. A cyclic constraint is adopted to regularize the deformation trajectories in conjunction with an enforcement of non-negative cortical thickness to ensure biological plausibility.

### 3.1 Coupled Learning of Cortical Surfaces

The human cerebral cortex is a highly folded 2D sheet with peaks (i.e., gyri) and grooves (i.e., sulci) (Fischl, 2012). The distance between sulci is relatively larger on the inner surface than on the outer surface, and this simpler folding pattern on the inner surface facilitates better (visual) discrimination and surface tessellation topology (Dale et al., 1999). Moreover, our preliminary experiments have shown that the closer the initial surface is to its target surface, the higher the reconstruction accuracy. Thus, we propose using the *midthickness* layer, positioned halfway between the inner and outer surfaces, as a starting point to couple the reconstructions of both the inner and outer cortical surfaces, with three key advantages: 1) It establishes a one-to-one mapping by directly encoding the correspondence between the inner and outer surfaces, facilitating accurate cortical surface reconstruction and cortical thickness estimation. 2) It reduces the difficulty of learning “large” deformations, striking a balance between optimizing the inner and outer surfaces. 3) It alleviates the challenge of obtaining a topologically correct initialization midthickness surface, as it is less convoluted in deep sulci than the pial surface.

The CSR problem is solved by learning diffeomorphic deformations that deform the initialization midthickness surface 𝒮_0_ towards its target WM and pial surfaces, 𝒮_*W*_ and 𝒮_*G*_ (see Fig. 1). Taking into account the discrepancy between the initialization surface, 𝒮_0_, and the *true* midthickness surfaces, 𝒮_*M*_, our method also learns a diffeomorphic deformation to optimize the initialization midthickness surface. Specifically, our method learns a neural network to model three diffeomorphic flows jointly:

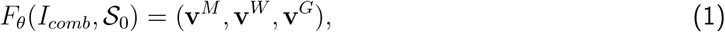

where *I*_*comb*_ is a multi-channel input consisting of brain MRI, cortical ribbon masks, and a signed distance function (SDF), and 𝒮_0_ is the initialization midthickness surface. The neural netowrk learns a set of three velocity fields to generate deformation fields: **v**^*M*^ generatesΦ_*M*_ to deform 𝒮_0_ to the *true* midthickness surface 𝒮_*M*_, **v**^*W*^ generatesΦ_*W*_ to deform 𝒮_*M*_ inward towards the WM surface 𝒮_*W*_, and **v**^*G*^ generatesΦ_*G*_ to deform 𝒮_*M*_ outward towards the pial surface 𝒮_*G*_. The joint learning establishs a *one-to-one correspondence* among 𝒮_*W*_, 𝒮_*M*_, and 𝒮_*G*_ to *explicitly couple* their joint reconstruction.

### 3.2 Loss Functions

Complementary loss functions are developed to both optimize the geometric precision of the reconstructed surfaces and regularize the velocity fields for diffeomorphic deformations.

#### Symmetric cycle loss

To enhance the diffeomorphism of the deformationsΦ_*W*_ andΦ_*G*_, which are modeled by separate velocity fields, we leverage the invertibility property of diffeomorphic mappings and introduce a symmetric cycle loss function during training to enforce deformation consistency of the midthickness surface along its entire trajectory and enhance their coupling. Fig. 2(a) demonstrates the deformation of a vertex **p**_*Mid*_ outward towards **p**_*GM*_ using **v**^*G*^ followed by a deformation inward towards 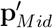 using **v**^*W*^, aiming to align two trajectories by minimizing the distance between **p**_*Mid*_ and 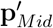.

**Figure 2:**
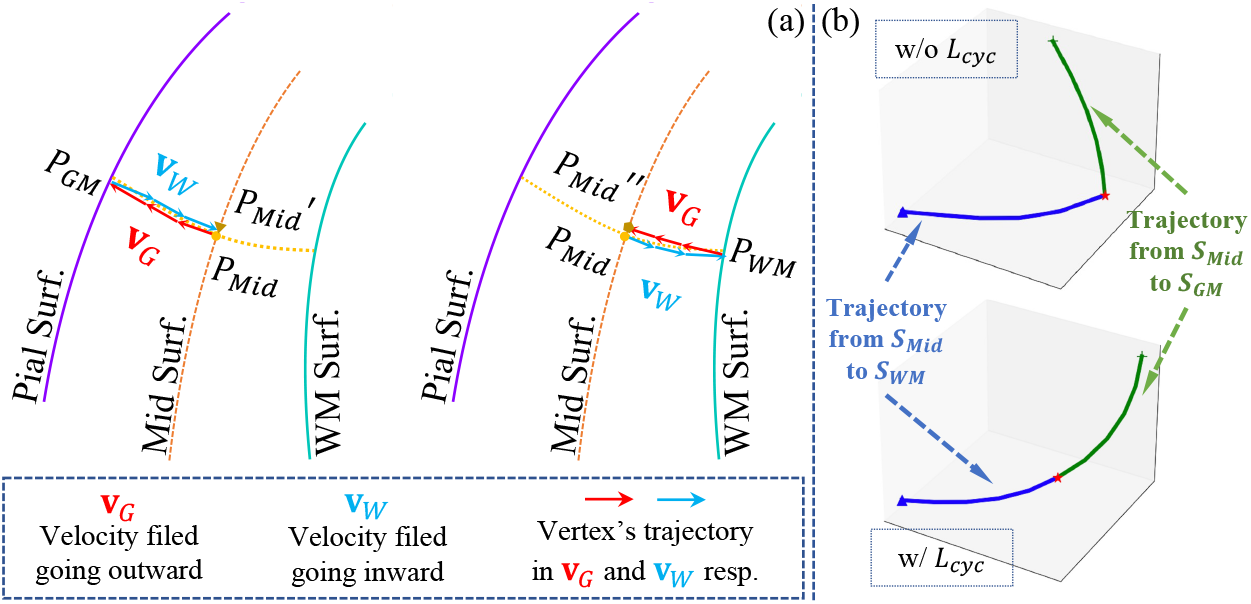
(a) Inverse consistency in coupled cortical surface reconstruction. Left: Deform a vertex **p**_*Mid*_ outward and inward towards 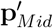 sequentially; Right: Deform **p**_*Mid*_ inward and outward towards 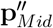 sequentially. (b) Example vertex deformation trajectories with and without ℒ_*cyc*_.

Similarly, 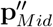 should be as close as possible to **p**_*Mid*_ when deformed inward byΦ_*W*_ and then outward byΦ_*G*_. The symmetric cycle loss ℒ_*cyc*_ is formulated as:

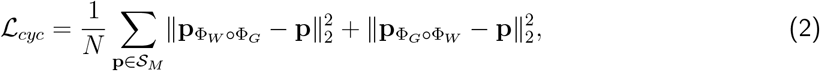

where 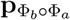 denotes the deformation of vertex **p** ∈ 𝒮_*M*_, first by velocity field **v**^a^ and then by **v**^b^ .

The cycle loss enhances the diffeomorphism of the deformation fields by ensuring the alignment of 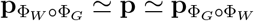,facilitating effective coupling of multiple diffeomorphic flows during the network optimization, as illustrated in Fig. 2(b). This regularization term also enables tracing the deformation trajectory of each vertex, facilitating vertex-wise cortical thickness estimation.

#### Mesh loss

It aims to minimize the distances of the vertices between the predicted surface meshes 𝒮_*W*_ (and 𝒮_*G*_) and their corresponding ground truth (GT) meshes 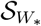 (and 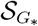) by the bidirectional Chamfer distance (Lebrat et al., 2021):

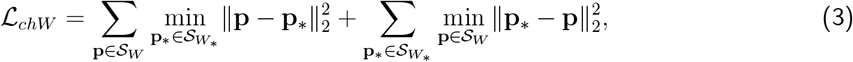

where **p** and **p**_*_ represent the vertex coordinates on the predicted and GT meshes, respectively. The loss for the pial surface, ℒ_*chG*_, is computed analogously. The total mesh loss combines the losses for both reconstructed surfaces: ℒ_*ch*_ = ℒ_*chW*_ + ℒ_*chG*_.

#### Trajectory loss

Starting from the midthickness layer, the trajectory length of the vertex moving towards the WM and the pial surfaces should be equal. We define the trajectory loss as the mean square difference of the trajectory lengths for each vertex:

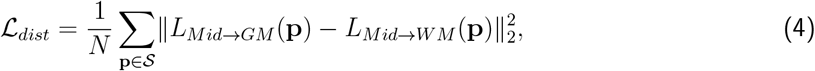

where 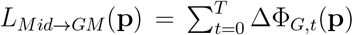 represents the accumulated trajectory length over *T* steps of deformation.

#### Normal consistency loss

We also employ a normal consistency regularization term to promote the surfaces’ smoothness:

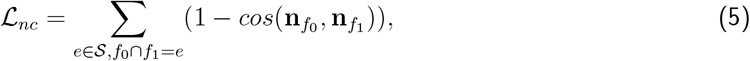

where *e* represents an edge, and *f*_0_ and *f*_1_ are are the two neighboring faces sharing *e*, with their unit normals denoted as 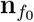 and 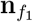,respectively.

We combine all the loss terms to optimize the model:

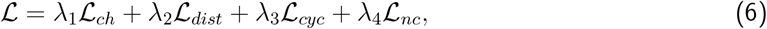

where {*λ*_*i*_}_*i*=1,···,4_ are weights to balance the loss terms. We empirically set *λ*_*i*_ = 1 for *i* = 1, · · ·, 3, and *λ*_4_ = 0.001.

### 3.3 Diffeomorphic Deformation Formulation

We aim to establish vertex-level correspondence between the initialization surface and the target surfaces while preserving the desired topology using diffeomorphic flows. Specifically, the diffeomorphic mapping is realized as the integration of a regularized velocity field **v**: Ω ⊂ ℝ^3^ × [0, 1] 1→ Ω ⊂ ℝ^3^ (denoted asΦ(**x**, *t*)), governed by an ODE (Arnold, 1992):

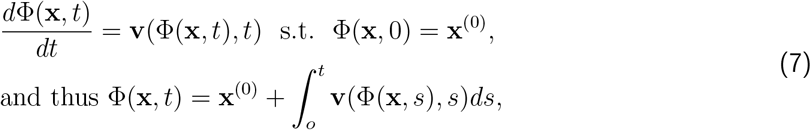

whereΦ(**x**, *t*) defines a trajectory from the source position **x**^(0)^ =Φ(**x**, 0) to the target position **x**^(1)^ =Φ(**x**, 1). According to the *Cauchy-Lipschitz* theorem (Teschl, 2012), if the velocity field is Lipschitz continuous, the resulting mappingΦ is bijective and possesses a continuous inverse (i.e., diffeomorphism). To solve this initial value problem (IVP), we employ a neural network to model the velocity fields and diffeomorphic deformation modules (DDMs) to compute the deformations, as illustrated in Fig. 1. Specifically, the DDM performs the integration by standard numerical integration techniques, such as the Euler method and the Runge-Kutta method (Burden et al., 2015).

The velocity fields can be modeled as either volume-wise or vertex-wise flows, referred to as the *dense-VF-based* method (SurfNet_D_) and the *vertex-VF-based* method (SurfNet_V_), respectively. Details are provided in subsections 3.4.3 and 3.4.4.

### 3.4 Framework Implementation

#### 3.4.1 Midthickness Surface Initialization

Existing CSR methods typically adopt an initialization surface that is either an oversmoothed template surface estimated from a group of subjects (Bongratz et al., 2022; Lebrat et al., 2021) or a WM surface (Q. Ma et al., 2021, 2022). While it is straightforward to initialize the midthickness surface by inflating the WM surface along its normal directions with a fixed predetermined distance, this approach is suboptimal because the thickness distribution is not uniform across the brain and the normal direction is merely an approximation based on neighboring faces. As shown in Fig. 3, we initialize the midthickness surface by leveraging the *local* WM and GM structures in the cortex ribbon segmentation maps (i.e., the filled interior area between WM and pial surfaces), resulting in a more accurate approximation of the true midthickness surface.

**Figure 3:**
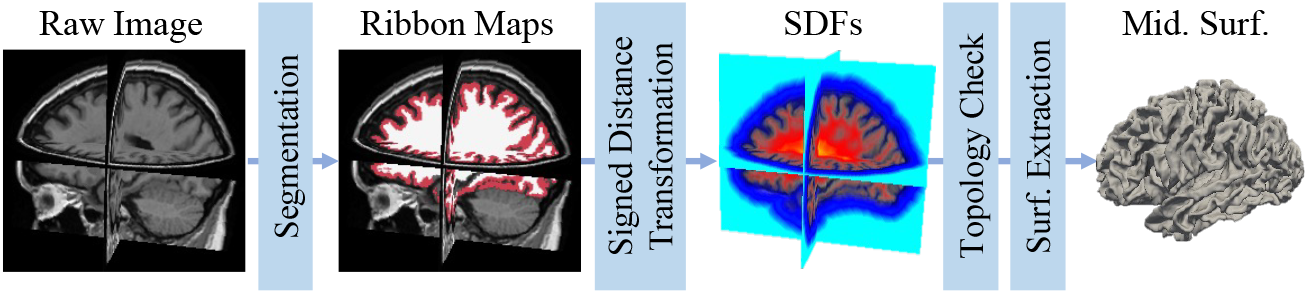
Midthickness surface initialization.

First, we train a 3D U-Net (Ronneberger et al., 2015) on a large-scale public neuroimaging dataset (Jack Jr et al., 2008) to generate the WM and GM segmentation maps from an input brain MRI volume. Second, we generate a signed distance function (SDF), *K*_*W*_, for the WM surface by using a distance transform algorithm, where voxels with zero values represent the surface boundaries and voxels with negative or positive values encode their distances to the surface boundaries inward or outward, respectively. A similar SDF, *K*_*G*_, is generated for the pial surface. The SDF whose 0-level implicitly defines the midthickness surface is obtained by averaging the WM and GM SDFs: *K*_*M*_ = (*K*_*G*_ + *K*_*W*_)*/*2. Third, we apply a fast topology check and correction algorithm (Bazin & Pham, 2007) to *K*_*M*_ to ensure the midthickness surface maintains spherical topology, in line with an existing study (Q. Ma et al., 2022). Finally, the initialization midthickness surface is extracted by the Marching Cubes algorithm (Lewiner et al., 2003) from the 0-level of *K*_*M*_ and parameterized by a triangular mesh 𝒮_0_.

#### 3.4.2 Feature Extraction

DL-based CSR studies have demonstrated the importance of integrating image features derived from MRI scans using CNNs and geometric features obtained from surface meshes through GNNs or FC-Nets. However, these methods typically extract the image and geometry features *separately* before information fusion, which limits the ability to *synergically* learn from heterogeneous features. To address this issue, we proposed to extract informative features from a multi-channel input consisting of a brain MRI, its cortical ribbon segmentation maps, and its midthickness surface represented as an SDF: *I*_*comb*_ = *I*©*M*_*W* ⊕*G*_©*K*_*M*_, where © is channel-wise concatenation and *M*_*W* ⊕*G*_ is a multi-class mask (BG=0, WM=0.5, GM=1).

This approach offers two main advantages: (1) Integrating heterogeneous information facilitates mutual knowledge distillation. While brain MRI provides intricate texture and semantics, it also contains noise and irrelevant regions far from the target surfaces. The cortical ribbon segmentation maps offer structural cues, aiding feature extraction near the surface boundaries. The SDF encapsulates the surface locations and spatial relationships between all voxels. The combination of these inputs offers a richer and more complementary learning source. (2) Employing a shared convolutional encoder for the entire 3D volume improves efficiency. Features are extracted in a single forward pass for all coordinates, obviating the need to sample local cubes and learn features independently. As mesh granularity increases, efficient interpolation within the feature space becomes viable.

#### 3.4.3 SurfNet_D_

The dense SVFs (**v**^*M*^, **v**^*W*^, **v**^*G*^) are parallel, separate predictions from the last layer of the 3D FCN, and the parameters are shared within the FCN. While scaling and squaring (SS) method (Arsigny et al., 2006) can be applied to the rigid grid of SVFs, followed by trilinear interpolation (Intp(·)) to obtain the deformations for vertices in continuous coordinates, there are three limitations for this straightforward solution: (1) Interpolation on the grid of deformation field cannot guarantee the invertibility of diffeomorphic mapping. (2) Numerical errors are amplified in the two sequential steps of SS+Intp(·). (3) Computation on grid voxels off the surface vertices is unnecessary. Hence, following (Lebrat et al., 2021; Santa Cruz et al., 2022), we compute the integration vertex-wisely. As shown in Fig. 4(a), we obtain the velocity vector for a vertex **x** by interpolating its neighboring velocity vectors (**v**_N(**x**)_): **v**_**x**_ = Intp(**v**_N(**x**)_). The vertex then moves to a new coordinate with a step of deformation 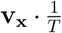,where *T* is the total time steps. The overall deformation can be obtained by repeating this procedure for *T* steps. Analogous to the proof in (Lebrat et al., 2021), Lipschitz continuity of the velocity field ensures the existence and uniqueness of the solution to Eq. 7. In numerical approximations of the solution, a sufficiently large integration time *T* in the ODE solver guarantees a diffeomorphism.

**Figure 4:**
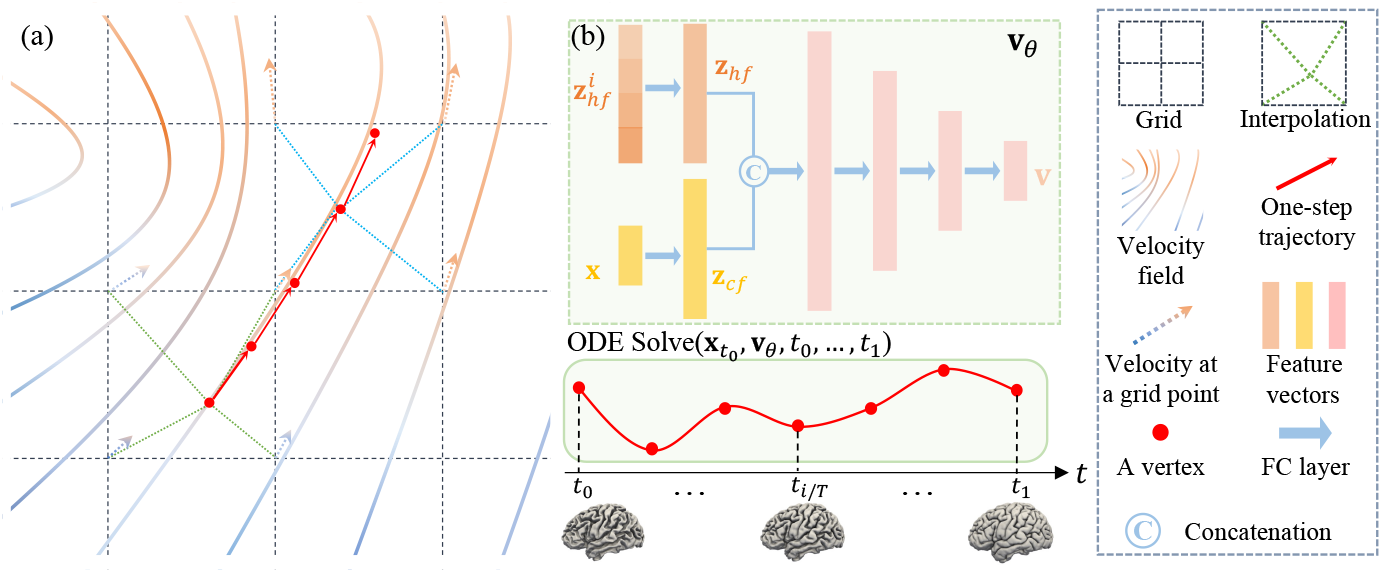
(a) The DDM in SurfNet_D_. (b) The DDM in SurfNet_V_ combines heterogeneous features with coordinate information to model the vertex’s trajectory by a NODE.

#### 3.4.4 SurfNet_V_

To parameterize the derivatives of the hidden state of each vertex using a NODE (R. T. Chen et al., 2018), we use a shared convolutional encoder and an FC-Net to parameterize the VF, and devise the *conditional* DDM to learn the deformation:

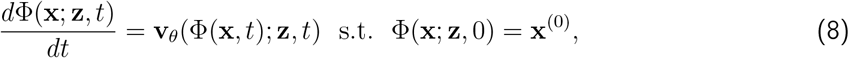

where **z** = {**z**_*hf*_, **z**_*cf*_ } is the conditional feature embedding extracted from the input image by a shared encoder and the surface by FC-layers respectively.

As shown in Fig. 4(b), the DDM in SurfNet_V_ is a Y-shape fully-connected network (FC-Net), inspired by previous NODEs-based works (Gupta & Chandraker, 2020; Q. Ma et al., 2022; Sun et al., 2022). The upper branch encodes heterogeneous features 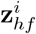’s by an FC layer: 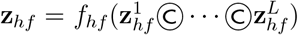,where © is channel-wise concatenation. Specifically, we employ a shared encoder with four scales to sequentially downsample the multi-channel input image *I*_*comb*_, forming a feature pyramid, 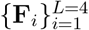. We then extract the 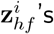 from the feature pyramid using the current surface vertices’ coordinates: 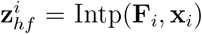, where **x**_*i*_ is normalized coordinates at the corresponding scale. With each deformation step, 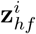 also updates according to the updated coordinates **x**_*i*_. The lower branch encodes the coordinate feature **z**_*cf*_ from each vertex’s current location by an FC layer: **z**_*cf*_ = *f*_*cf*_ (**x**), which serves as complementary information for each vertex. Then **z**_*hf*_ and **z**_*cf*_ are concatenated and followed by four FC layers, each followed by a non-linear activation function ReLU. Particularly, besides the shared encoder, each DDM has its distinct parameters that are not shared, i.e., 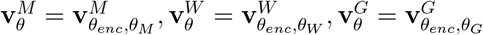. . Analogous to the proof in (Q. Ma et al., 2022), the Lipschitz continuity of the deformation network theoretically prevents surface self-intersections. In practice, a larger value of *T* further reduces surface intersections introduced by the numerical approximation in ODE discretization while maintaining constant memory cost (R. T. Chen et al., 2018; Lebrat et al., 2021; Q. Ma et al., 2022).

#### 3.4.5 Network Training

We utilize the loss function of Eq. 6 to supervise the network training in an *end-to-end* manner. For SurfNet_D_, the DDM is an ODE solver with no trainable parameters, and the whole network is optimized by standard back-propagation. For SurfNet_V_, the DDM is parameterized by the NODE (Fig. 4(b)), and gradients are computed using the *adjoint sensitivity method* (R. T. Chen et al., 2018).

## 4 Experiments

We have evaluated our method for reconstructing both WM and pial surfaces on large-scale datasets, including (ADNI) (Jack Jr et al., 2008), HCP (Van Essen et al., 2013), and OASIS (Marcus et al., 2007).

### 4.1 Experimental Setups

#### 4.1.1 Datasets

The ADNI-1 (Jack Jr et al., 2008) dataset consists of 817 subjects aged 55 to 90. We randomly split it into subsets of 654, 50, and 113 subjects for training, validation, and testing, respectively. The HCP (Van Essen et al., 2013) dataset consists of 1113 subjects aged 22 to 35. We randomly chose 890, 75, and 148 subjects for training, validation, and testing, respectively. The OASIS-1 (Marcus et al., 2007) dataset consists of 413 subjects aged 18 to 96. We randomly split it into subsets of 330, 25, and 58 subjects for training, validation, and testing, respectively. The models were trained on the training sets until they reached a loss plateau on the validation set, after which their performance was evaluated on the test sets. We followed a pre-processing protocol used in existing studies (Bongratz et al., 2022; Cruz et al., 2021; Lebrat et al., 2021; Q. Ma et al., 2022) for fair comparison. The T1-weighted MRI scans were aligned to the MNI152 template and clipped to have the size of 192 × 224 × 192 at 1*mm*^3^ isotropic resolution. The pseudo ground-truth (GT) of ribbon segmentation and cortical surfaces were generated using FreeSurfer v7.2.0 (Fischl, 2012). The intensity values of MRI scans, ribbon segmentation maps, and SDFs were normalized to [0, 1] and the coordinates of the vertices were normalized to [−1, 1].

#### 4.1.2 Implementation Details

Our framework was implemented in PyTorch (Paszke et al., 2019) and trained on a workstation with 12 GB NVIDIA P100 GPU. The 3D U-Net (Ronneberger et al., 2015) for segmenting ribbons was trained for 200 epochs using Adam (Kingma & Ba, 2015) optimization and achieved an average Dice index of 0.96 on the testing set. The SurfNet model utilized *T* = 10 steps (i.e., step size is 0.1) in Euler solver. We trained SurfNet model using Adam optimizer (*β*_1_ = 0.9, *β*_2_ = 0.999, *ϵ* = 1*e*^−10^, learning rate = 1*e*^−4^) for 400 epochs to reconstruct WM, midthickness, and pial surfaces of both brain hemispheres. The surface meshes had ∼130*k* vertices.

#### 4.1.3 Evaluation Metrics

We utilized three distance-based metrics to measure the CSR accuracy. The Chamfer Distance (CD) between two sets of points *A* and *B* is defined as the sum of squared distances from each point in set *A* to its nearest neighbor in set *B*, and vice versa (Fan et al., 2017; N. Wang et al., 2020):

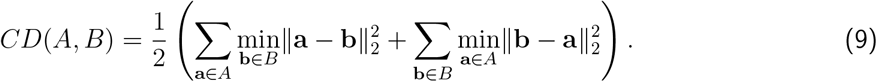

The Average Symmetric Surface Distance (ASSD) is the average of the distance measures from each point on the first surface to the nearest point on the second surface and vice versa (Cruz et al., 2021):

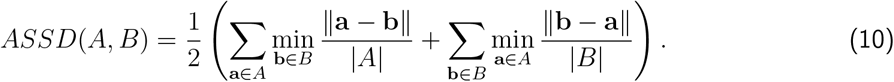

The Hausdorff distance (HD) measures the maximum distance of a point set to the nearest point in the other set (Cruz et al., 2021; Taha & Hanbury, 2015). We used the 90th percentile instead of the maximum because HD is sensitive to outliers (Huttenlocher et al., 1993):

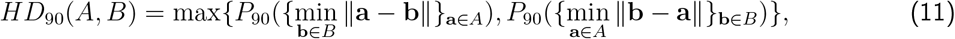

where *P*_90_({*d*_*i*_}) denotes the 90th percentile of the set of distances {*d*_*i*_}. They were computed bidirectionally over ∼130*k* points uniformly sampled from the predicted and target surfaces. A lower distance means a better result. Since topology is also important in CSR, we utilized the ratio of self-intersection faces (SIF) (Cruz et al., 2021; Dahnke et al., 2013; Q. Ma et al., 2022) to measure quality of the reconstructed surfaces.

#### 4.1.5 Baselines

We compared SurfNet with representatives from the two categories of existing DL-based CSR methods and evaluated their performance for the reconstruction of both WM and pial surfaces. Particularly, DeepCSR (Cruz et al., 2021) represents implicit surface reconstruction methods, while others fall into the category of explicit methods. PialNN (Q. Ma et al., 2021) is designed for pial surface reconstruction by deforming an initial WM surface. CorticalFlow (Lebrat et al., 2021) utilizes convex hulls as the initialization template with the first-order Euler solver, whereas CorticalFlow++ (Santa Cruz et al., 2022) utilizes smoothed convex hulls and the fourth-order Runge-Kutta (RK4) solver. *Surf* NN (Zheng et al., 2023b) initializes a subject-specific midthickness surface, learns deformation through a CNN-based network, and predicts the half cortical thickness. ccfCSR (Dong et al., 2024) gradually deforms a surface mesh template towards the inner surface and subsequently towards the pial surface, followed by iterative cyclical refinement until convergence. CortexODE (Q. Ma et al., 2022) employs WM segmentation for surface initialization and NODE for deformation computation. Vox2cortex (Bongratz et al., 2022) deforms averaged surface templates with a GNN-based network to reconstruct multiple surfaces.

### 4.2 Comparison with SOTA CSR Methods

#### 4.2.1 Main Results & Analysis

Our method achieved substantial improvement on both the WM and pial surfaces over other approaches. Since DeepCSR (Cruz et al., 2021) predicts a SDF based implicit surface and requires post-processing to correct topology and extract a mesh, its results, while potentially free of SIFs, were less accurate compared with those of the explicit CSR methods. Starting from a WM surface, PialNN (Q. Ma et al., 2021) can achieve sub-voxel accuracy for reconstructing the pial surface but its SIF ratio was relatively high. Vox2Cortex (Q. Ma et al., 2022) can generate multiple surfaces from different template meshes. It employs complex CNN and GNN models to model the deformation for each vertex but has no diffeomorphism guarantee. The promising results of (Dong et al., 2024; Lebrat et al., 2021; Q. Ma et al., 2022; Santa Cruz et al., 2022) indicated that using a neural network to parameterize the ODE can facilitate diffeomorphic deformation, yielding better CSR accuracy and better topological quality. However, methods that start from oversmoothed templates (Bongratz et al., 2022; Dong et al., 2024; Lebrat et al., 2021; Santa Cruz et al., 2022) generally yield inferior results compared to those utilizing finer initialization surfaces (Q. Ma et al., 2022; Zheng et al., 2023b). Our method achieved the overall best performance, due to its improved initialization surface, coupled reconstruction of multiple surfaces, and explicit regularizations of vortex deformation trajectories.

On the ADNI dataset, our method SurfNet_D_ achieved ∼51.2% improvement in mean ASSD (of WM and pial surfaces) compared to the best baseline CortexODE (i.e., 0.099*mm v*.*s* 0.203*mm*). Our method effectively reduced surface self-intersections measured by the percentage of self-intersecting faces (SIFs). SurfNet_V_ almost produced no SIFs on the WM surface and ∼0.013% SIFs on the pial surface, gaining an average improvement of more than 50% compared to CortexODE. Although SurfNet_D_ had higher SIFs than CortexODE, its performance was competitive (average ∼0.026% *v*.*s* ∼0.019%) and outperformed the baseline methods. Moreover, the reconstructed WM surfaces exhibited higher accuracy and superior surface quality. We speculate that this may attribute to the fact that the pial surface is more convoluted and has a smaller spatial distance between deep sulci than the WM surface, making it more challenging to deform the mid-thickness surface to the pial surface. On the OASIS and HCP datasets, our method achieved consistent performance improvements and surpassed the alternative methods.

#### 4.2.2 SurfNet_D_ v.s SurfNet_V_

As shown in Table 2, SurfNet_D_ consistently outperformed SurfNet_V_ by a margin in terms of the reconstructed surface accuracy (i.e., CD, ASSD, HD), while SurfNet_V_ exhibited lower SIFs on both WM and pial surfaces. This discrepancy can be attributed to differences in the implementations of the ODE solver and the optimization of the network. Specifically, SurfNet_D_ employs an FCN to predict dense 3D SVFs, samples vertex-wise velocity via interpolation, and numerically solve Eq. 7, while SurfNet_V_ samples features from extracted feature maps conditioned on vertex coordinates and adopts neural ODEs to solve Eq. 8. The former network was optimized by a standard back-propagation, while the latter computed gradients using the adjoint sensitivity method (R. T. Chen et al., 2018). In addtion to sufficiently small integration step size, SurfNet_D_’s performance depends on the smoothness of the SVFs, whereas SurfNet_V_’s performance relies on the Lipschitz continuity of the network.

**Table 2:** Quantitative analysis of cortical surface reconstruction on geometric accuracy and self-intersections. The Chamfer distance (CD), average symmetric surface distance (ASSD), Hausdorff distance (HD), and the ratio of the self-intersecting faces (SIF) were measured for WM and pial surfaces on three datasets. The mean value and standard deviation are reported. For all metrics, lower scores mean better results. The best and second-best ones are in **red** and **blue**, respectively.

|  | Method | L-Pial Surface |  |  |  | L-WM Surface |  |  |  |
| --- | --- | --- | --- | --- | --- | --- | --- | --- | --- |
|  |  | CD (mm) | ASSD (mm) | HD (mm) | SIF (%) | CD (mm) | ASSD (mm) | HD (mm) | SIF (%) |
| ADNI | DeepCSR (Cruz et al., 2021) | 0.945±0.078 | 0.593±0.065 | 1.149±0.203 | \ | 0.938±0.076 | 0.587±0.064 | 1.137±0.193 | \ |
|  | PialINN (Q. Ma et al., 2021) | 0.621±0.035 | 0.465±0.044 | 1.002±0.106 | 0.137±0.093 | \ | \ | \ | \ |
|  | CorticalFlow (Lebrat et al., 2021) | 0.691±0.043 | 0.497±0.049 | 1.106±0.115 | 0.149±0.087 | 0.641±0.037 | 0.465±0.042 | 0.996±0.100 | 0.108±0.073 |
|  | CorticalFlow++ (Santa Cruz et al., 2022) | 0.545±0.036 | 0.410±0.033 | 0.886±0.069 | 0.098±0.067 | 0.544±0.034 | 0.401±0.030 | 0.878±0.066 | 0.069±0.042 |
|  | Vox2Cortex (Bongratz et al., 2022) | 0.582±0.028 | 0.370±0.025 | 0.746±0.057 | 0.059±0.039 | 0.577±0.027 | 0.353±0.022 | 0.722±0.055 | 0.043±0.023 |
|  | SurfNN (Zheng et al., 2023b) | 0.402±0.021 | 0.228±0.022 | 0.518±0.061 | 0.168±0.117 | 0.343±0.018 | 0.165±0.014 | 0.368±0.032 | 0.136±0.089 |
|  | ccfCSR (Dong et al., 2024) | 0.496±0.040 | 0.242±0.039 | 0.623±0.057 | 0.052±0.043 | 0.490±0.037 | 0.239±0.033 | 0.609±0.051 | 0.039±0.030 |
|  | cortexODE (Q. Ma et al., 2022) | 0.476±0.017 | 0.214±0.020 | 0.455±0.058 | **0.022**±0.012 | 0.458±0.016 | 0.192±0.015 | 0.436±0.014 | **0.015**±0.011 |
|  | SurfNet<sub>V</sub> | **0.322**±0.021 | **0.123**±0.010 | **0.267**±0.022 | **0.013**±0.011 | **0.303**±0.018 | **0.117**±0.010 | **0.254**±0.021 | **0.005**±0.002 |
|  | SurfNet<sub>D</sub> | **0.310**±0.017 | **0.111**±0.009 | **0.250**±0.017 | 0.031±0.019 | **0.253**±0.015 | **0.087**±0.007 | **0.190**±0.015 | 0.021±0.014 |
| HCP | DeepCSR (Cruz et al., 2021) | 0.979±0.082 | 0.598±0.067 | 1.282±0.211 | \ | 0.944±0.079 | 0.589±0.064 | 1.141±0.194 | \ |
|  | PialINN (Q. Ma et al., 2021) | 0.630±0.029 | 0.476±0.041 | 1.022±0.108 | 0.142±0.094 | \ | \ | \ | \ |
|  | CorticalFlow (Lebrat et al., 2021) | 0.689±0.040 | 0.494±0.046 | 1.079±0.109 | 0.147±0.087 | 0.640±0.037 | 0.460±0.038 | 0.988±0.096 | 0.100±0.070 |
|  | CorticalFlow++ (Santa Cruz et al., 2022) | 0.534±0.037 | 0.402±0.031 | 0.811±0.057 | 0.086±0.043 | 0.467±0.020 | 0.399±0.030 | 0.813±0.058 | 0.086±0.040 |
|  | Vox2Cortex (Bongratz et al., 2022) | 0.587±0.020 | 0.378±0.026 | 0.749±0.058 | 0.060±0.043 | 0.578±0.029 | 0.357±0.023 | 0.726±0.056 | 0.039±0.021 |
|  | SurfNN (Zheng et al., 2023b) | 0.449±0.024 | 0.254±0.024 | 0.577±0.066 | 0.185±0.130 | 0.373±0.020 | 0.182±0.015 | 0.399±0.034 | 0.142±0.092 |
|  | ccfCSR (Dong et al., 2024) | 0.548±0.045 | 0.269±0.041 | 0.697±0.063 | 0.058±0.046 | 0.529±0.039 | 0.260±0.034 | 0.651±0.053 | 0.040±0.030 |
|  | cortexODE (Q. Ma et al., 2022) | 0.483±0.021 | 0.219±0.019 | 0.464±0.062 | **0.025**±0.014 | 0.461±0.017 | 0.203±0.014 | 0.437±0.014 | **0.018**±0.013 |
|  | SurfNet<sub>V</sub> | **0.400**±0.031 | **0.139**±0.016 | **0.278**±0.020 | **0.015**±0.012 | **0.362**±0.019 | **0.131**±0.022 | **0.277**±0.026 | **0.006**±0.002 |
|  | SurfNet<sub>D</sub> | **0.372**±0.015 | **0.120**±0.008 | **0.260**±0.016 | 0.035±0.022 | **0.304**±0.012 | **0.098**±0.007 | **0.211**±0.013 | 0.024±0.017 |
| OASIS | DeepCSR (Cruz et al., 2021) | 0.986±0.085 | 0.617±0.070 | 1.331±0.212 | \ | 0.975±0.081 | 0.594±0.067 | 1.151±0.197 | \ |
|  | PialINN (Q. Ma et al., 2021) | 0.635±0.032 | 0.460±0.038 | 0.993±0.082 | 0.141±0.096 | \ | \ | \ | \ |
|  | CorticalFlow (Lebrat et al., 2021) | 0.687±0.040 | 0.495±0.047 | 1.082±0.110 | 0.147±0.086 | 0.637±0.035 | 0.462±0.040 | 0.992±0.097 | 0.101±0.070 |
|  | CorticalFlow++ (Santa Cruz et al., 2022) | 0.531±0.035 | 0.399±0.030 | 0.812±0.057 | 0.088±0.045 | 0.529±0.033 | 0.398±0.030 | 0.810±0.055 | 0.086±0.042 |
|  | Vox2Cortex (Bongratz et al., 2022) | 0.588±0.032 | 0.381±0.030 | 0.750±0.063 | 0.061±0.037 | 0.581±0.028 | 0.375±0.027 | 0.731±0.059 | 0.046±0.027 |
|  | SurfNN (Zheng et al., 2023b) | 0.454±0.024 | 0.258±0.025 | 0.596±0.069 | 0.188±0.132 | 0.374±0.020 | 0.182±0.015 | 0.404±0.035 | 0.149±0.098 |
|  | ccfCSR (Dong et al., 2024) | 0.557±0.045 | 0.273±0.044 | 0.703±0.064 | 0.059±0.047 | 0.536±0.040 | 0.261±0.038 | 0.669±0.056 | 0.041±0.032 |
|  | cortexODE (Q. Ma et al., 2022) | 0.481±0.019 | 0.218±0.021 | 0.461±0.062 | **0.026**±0.015 | 0.463±0.018 | 0.207±0.017 | 0.435±0.015 | **0.018**±0.010 |
|  | SurfNet<sub>V</sub> | **0.410**±0.034 | **0.142**±0.016 | **0.281**±0.024 | **0.016**±0.012 | **0.349**±0.024 | **0.128**±0.019 | **0.266**±0.022 | **0.007**±0.002 |
|  | SurfNet<sub>D</sub> | **0.384**±0.018 | **0.125**±0.010 | **0.258**±0.018 | 0.033±0.021 | **0.308**±0.016 | **0.101**±0.008 | **0.213**±0.016 | **0.018**±0.012 |

#### 4.2.3 Qualitative Comparison

We visualize the reconstructed cortical surfaces with color maps decoding their geometric errors and overlay their projected boundary onto MRI slices in Fig. 5. The CSR results obtained by our method had uniformly smaller errors across the whole surface with reduced artifacts (e.g., self-intersection, over-smoothness, corruption), compared with those generated by the alternatives.

**Figure 5:**
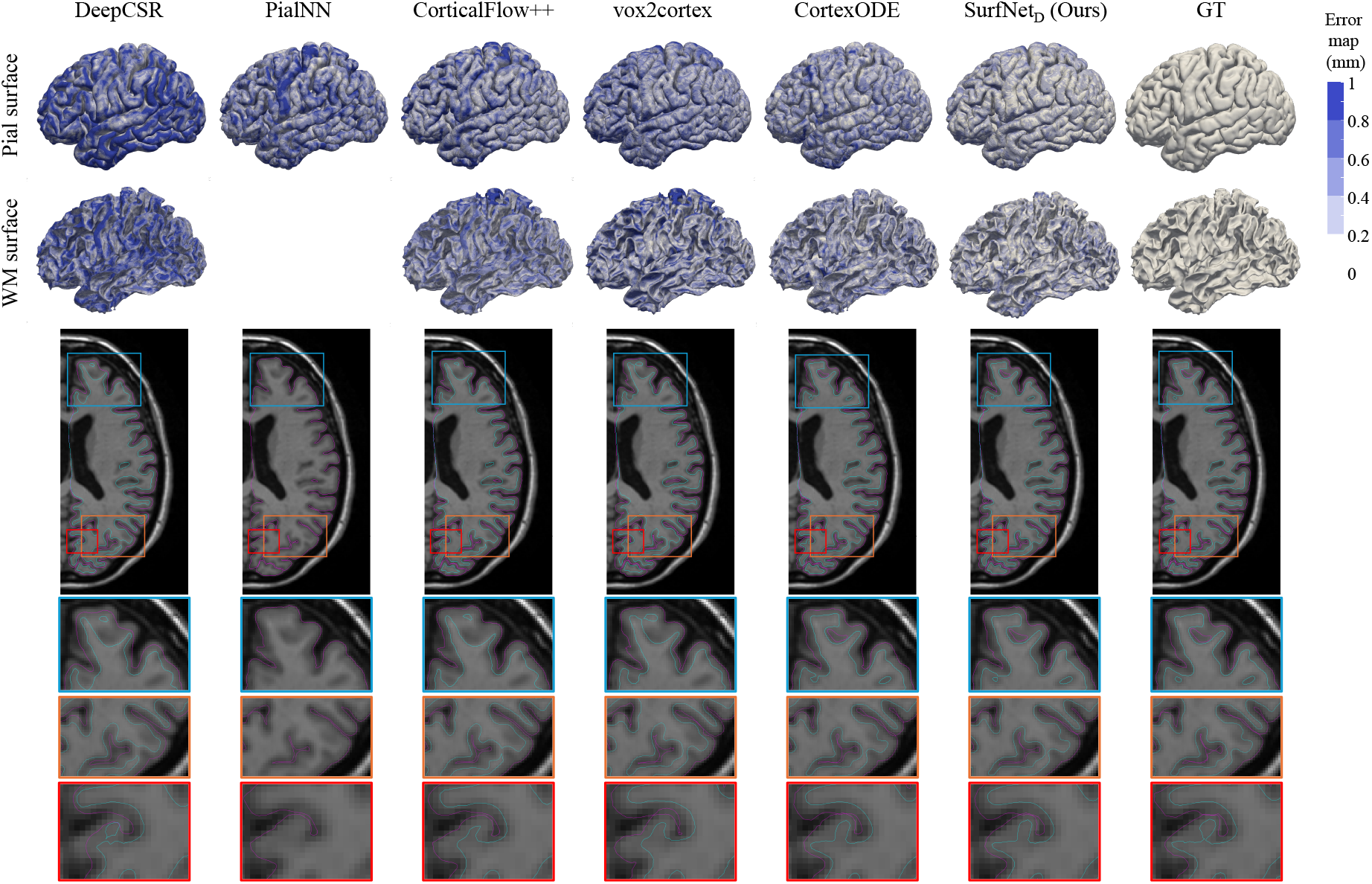
Top: Visualization of the reconstructed WM and pial surfaces with distance compared to the ground truth. Bottom: Visualization of WM surface (Cyan) and Pial surface (Magenta) overlaid on the MRI slice (best viewed in color).

#### 4.2.4 Computational Time

FreeSurfer requires 4∼6 hours to process a single subject, while DL-based CSR methods, including our SurfNet, are thousands of times faster. The inference time of all DL baselines is reported in Fig. 6. The runtime is measured on an NVIDIA P100 GPU. The DeepCSR framework requires 2 minutes due to the voxel-wise prediction on a high-resolution volume. CorticalFlow++ (Euler, 10 steps) achieves faster runtime because it has no volume-level processing. The SOTA explicit CSR method CortexODE (Q. Ma et al., 2022) needed ∼1*s* to reconstruct two surfaces *sequentially*. Our method, SurfNet_D_, simultaneously reconstructs *three* surfaces (i.e., WM, pial, and midthickness) in ∼0.24*s*. SurfNet_V_ requires more time for training and inference due to the higher computational cost of NODE, but its inference time is less than 1*s* for reconstructing three surfaces. Overall, our method is computationally as efficient as the SOTA alternatives and significantly faster than conventional FreeSurfer pipelines.

**Figure 6:**
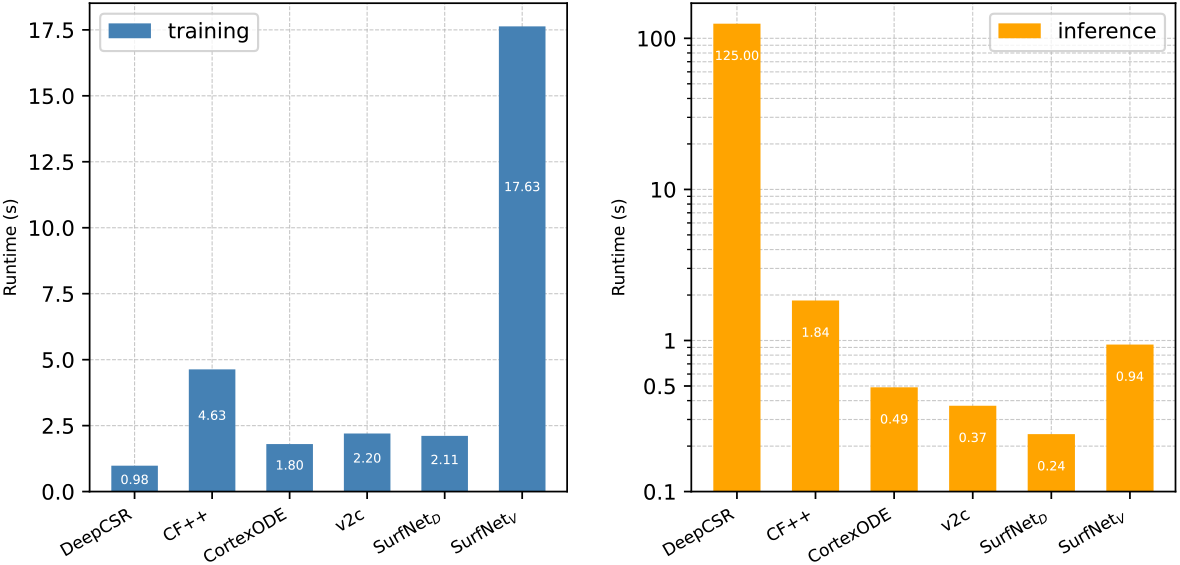
The runtime (in seconds) of one iteration of the neural network for the ADNI dataset during the training and inference stages, respectively.

### 4.3 Ablation Studies

We conducted ablation experiments based on SurfNet_D_ to verify the effectiveness of the components of our method.

#### 4.3.1 Input

Our method takes a multi-channel input of 3D images *I*_*comb*_ and an initialization midthickness surface *S*_0_. We conducted two experiments to analyze the influence of different components on the CSR reconstruction performance (Table 3 Top). *First*, by gradually removing components from *I*_*comb*_, we observed a significant drop in accuracy, indicating the contribution of both SDF and segmentation maps to the final results. *Second*, we investigated the impact of the initialization surface by applying Laplacian smoothing on *S*_0_ to generate a smoothed surface. The results revealed that the increased smoothness of the initialization surface decreased the model’s accuracy.

**Table 3:**
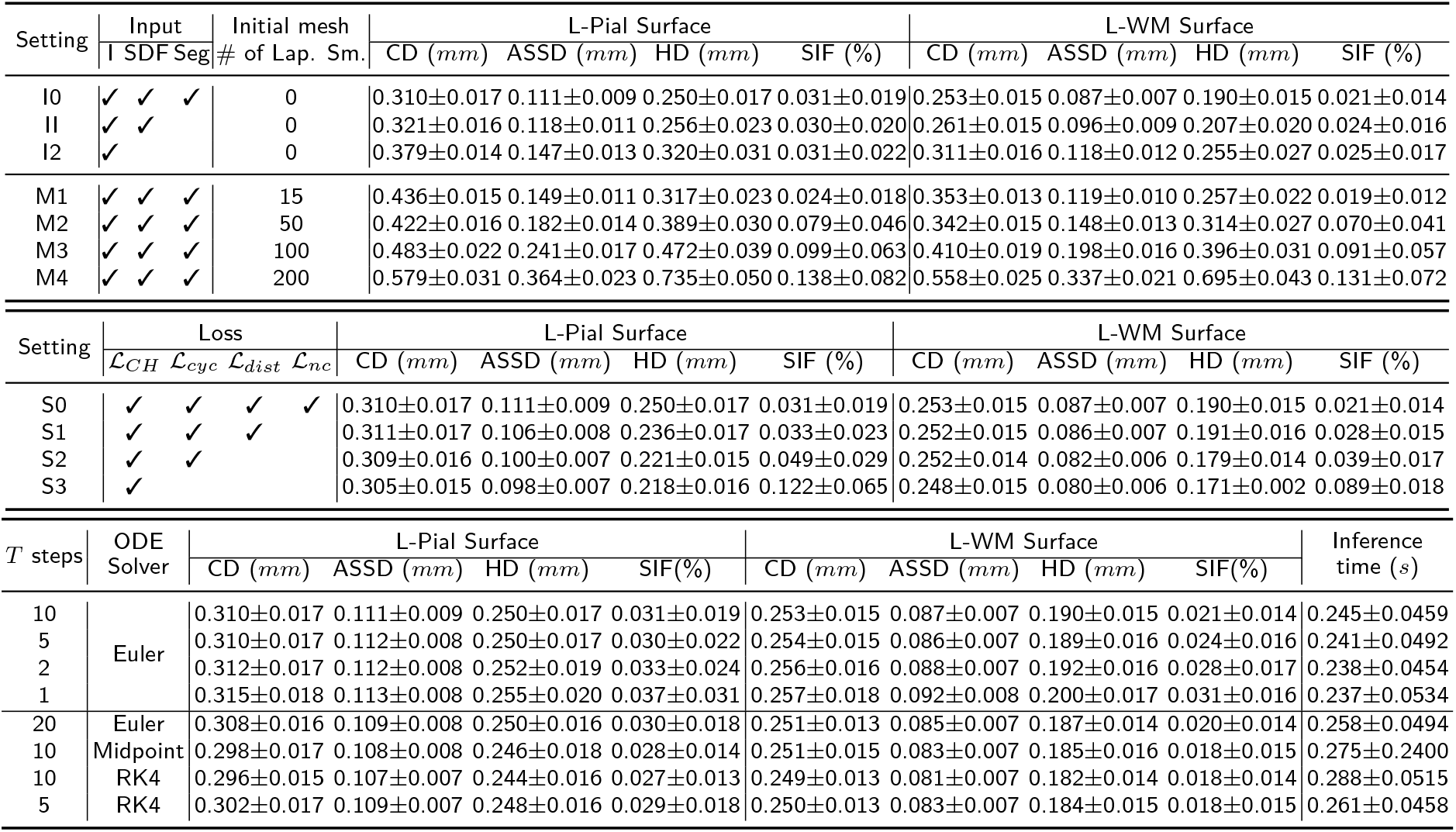
Ablation studies of SurfNet_D_ model on ADNI-1 dataset in terms of CD, ASSD, HD, and SIF. Top: The impact of the input data and initialization surface. Middle: The impact of loss functions. Bottom: The impact of deformation steps in diffeomorphic deformation module (DDM).

#### 4.3.2 Loss Functions

We evaluated various combinations of loss functions to train the model and summarized the results in Table 3 Middle. Particularly, the first row (referred to as S0) corresponds to our complete network setting, while the last row (S3) represents using Chamfer distance alone. Through the ablation studies, we observed that each component played its own role in a complementary way. The proposed symmetric cycle loss (S2, ℒ_*cyc*_) promoted the invertibility of deformations, yielding a significant reduction of self-intersections on the meshes. We visualize a sample vertex deformation trajectory in Fig. 2 to illustrate the difference between the settings of S2 and S3. Besides, enforcing equality of the trajectories from the midthickness surface to the WM and pial surfaces (S1, ℒ_*dist*_) helped optimize the midthickness surface, thereby preventing deformation in an arbitrary direction and reducing self-intersection. However, the geometric accuracy slightly decreased, which may caused by the difficulty in accessing highly curved regions or deep sulci under such strong topology constraints. Moreover, adopting smoothness regularization terms on the surfaces further enhance the surface quality (S0, ℒ_*nc*_).

We conducted a series of experiments to evaluate the sensitivity of the weights used to balance the loss terms, with results for both the WM and pial surfaces shown in Fig. 7. With *λ*_1_ (the weight for the mesh loss) fixed at 1, the mesh accuracy measured by CD showed slight degradation but remained relatively stable when *λ*_2_ (trajectory loss) and *λ*_3_ (symmetric cycle loss) were within the range of [0.75, 2]. Additionally, surface quality measured by SIF improved steadily as *λ*_2_ and *λ*_3_ increased from 0.25 to 2 and saturated for values above 1. As *λ*_4_ (normal consistency loss) increased from 1e-4 to 0.1, mesh accuracy decreased while mesh quality improved. The purpose of these experiments was not to identify the parameter values yielding the highest accuracy but to demonstrate that the selected weights fall within a range where a balance between geometric accuracy and topological quality is achieved, ensuring stable and high overall performance.

**Figure 7:**
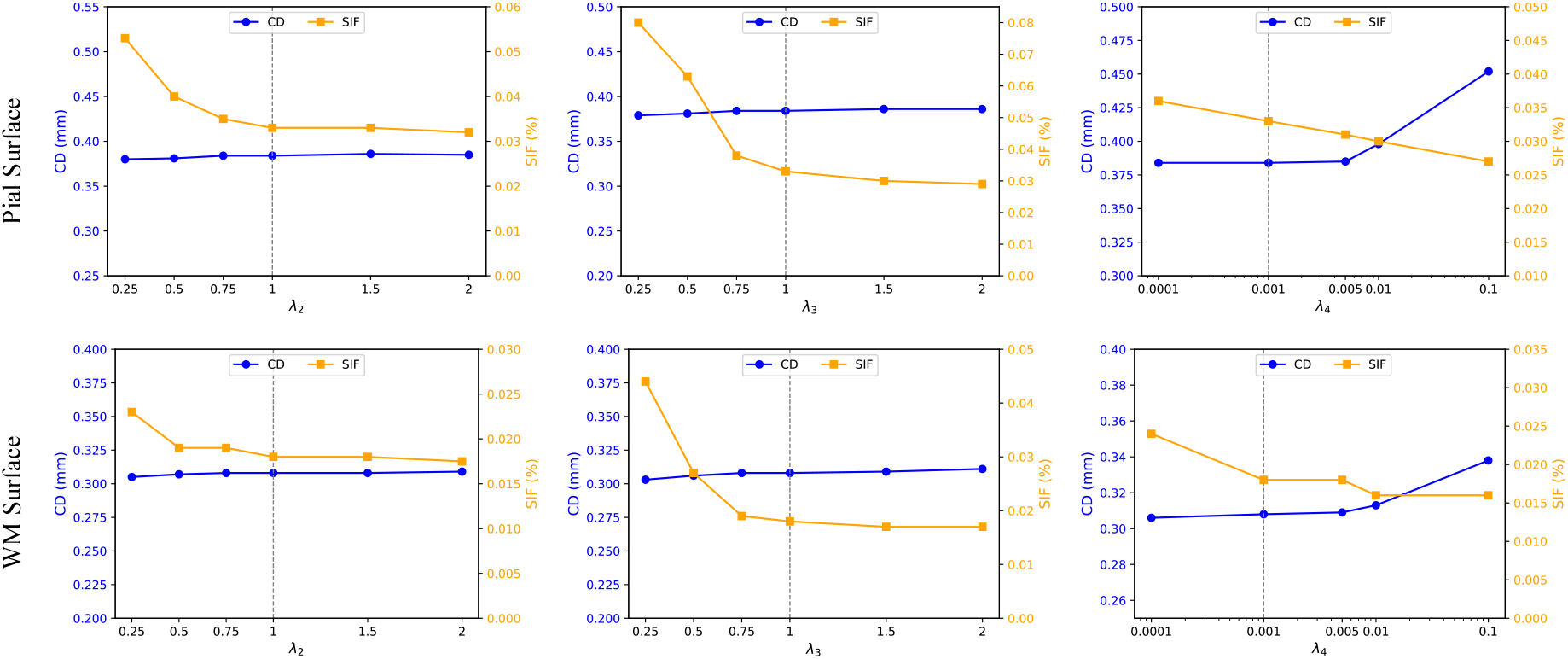
Evaluation of the stability of weights for balancing four loss terms. With *λ*_1_ (the weight for the mesh’s Chamfer distance) is fixed at 1, we assess the performance of SurfNet_D_ on OASIS dataset across three experiments by varying the weights, *λ*_2_ (trajectory loss), *λ*_3_ (symmetric cycle loss), and *λ*_4_ (normal consistency loss).

#### 4.3.3 ODE Solver in DDM

Table 3 Bottom summarizes evaluation results of the DDM computation in SurfNet_D_. *First*, we investigated the effect of different numbers of deformation steps (*T*) in DDM. The results revealed that as *T* increased, the performance first improved and then saturated, indicating that five to ten steps were sufficiently large to deform the midthickness surface to the inner or outer surfaces. *Second*, we explored different ODE solvers in DDM. Maintaining the same number of integration steps, employing more sophisticated numerical integration led to additional performance improvement, albeit at a cost of increased computational time. For instance, when *T* = 10, the ASSD improved by 2.7%, accompanied by a 12.2% increase in inference time for the Midpoint ODE solver. To strike a balance of efficiency and efficacy, we used Euler solver with *T* = 10 in our experiments.

#### 4.3.4 Ribbon Segmentation

Our method utilize ribbon segmentation maps to initialize the midthickness surface. To investigate how different segmentation results impact the final CSR performance, we use the ribbon maps and cortical surfaces generated by FreeSurfer (Fischl, 2012) pipeline as ground truth for comparison. The Dice coefficient of the predicted segmentation and the errors (ASSA, HD, SIFs) of the reconstructed cortical surfaces using different segmentation results are reported in Table 4. 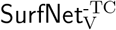 uses the predicted ribbon segmentation maps of our DL-based segmentation model without performing topology correction on the SDFs. This resulted in similar geometric errors in the reconstructed surfaces but significantly worse SIF, showing the necessity of topology checks and corrections in the midthickness surface initialization.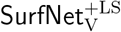 employs 100 iterations of Laplacian smoothing on our extracted midthickness surface, resulting in over-smoothed initialization surface that is topologically correct but geometrically far from the true midthickness surface. It led to much worse reconstruction accuracy and the large deformation led to worse SIF. Utilizing FreeSurfer’s segmentation maps led to the best performance, indicating that better CSR can be achieved with more accurate ribbon segmentation.

**Table 4:**
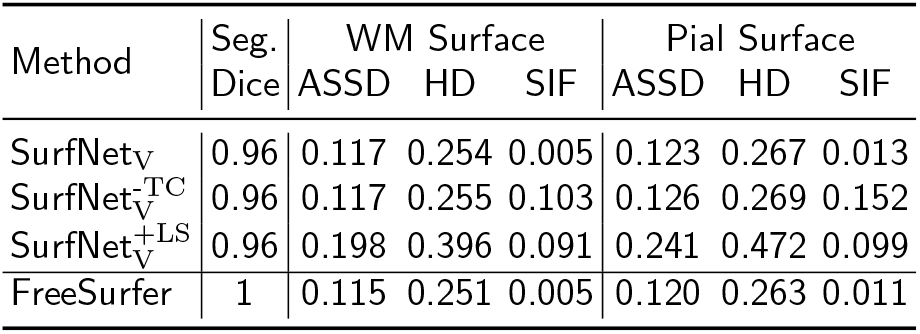
Impact of ribbon segmentation in CSR of SurfNet_V_ on the ADNI dataset. The Dice score, ASSA (*mm*), HD (*mm*), and SIFs (%) are compared with the GT from FreeSurfer.

### 4.4 Clinical Applications

#### 4.4.1 Reproducibility

The reproducibility of CSR methods measures the consistency of their outputs, which is crucial for clinical applications. We carried out reproducibility experiments on two datasets: a paired ADNI_1.5&3T_ dataset (Jack Jr et al., 2008) consisting of 1.5T and 3T MRIs of the same subjects, and the TestRetest dataset (Maclaren et al., 2014) comprising 40 MRIs collected within a short period for each of the 3 subjects. In these scenarios, the cortical surfaces of the same subject should be nearly identical. Following the experimental setup in prior studies (Bongratz et al., 2022; Cruz et al., 2021; Q. Ma et al., 2022; Zheng et al., 2023b), we utilized the iterative closest-point algorithm (ICP) to align each pair of images and computed the geometric distance between the corresponding surfaces. The results for the left WM surfaces are presented in Table 5, demonstrating that our method obtained superior reproducibility compared with FreeSurfer and was comparable to the SOTA DL methods.

**Table 5:**
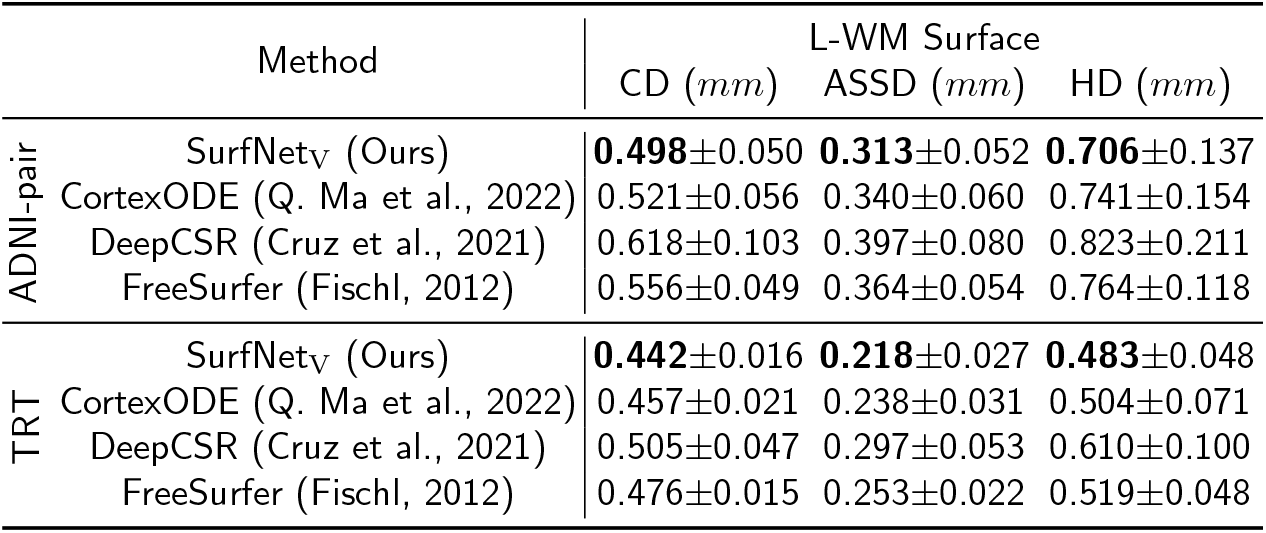
Reproducibility analysis.

|  | Method | L-WM Surface |  |  |
| --- | --- | --- | --- | --- |
| | | CD ($mm$) | ASSD ($mm$) | HD ($mm$) |
| ADNI-pair | SurfNet<sub>V</sub> (Ours) | **0.498**±0.050 | **0.313**±0.052 | **0.706**±0.137 |
| | CortexODE (Q. Ma et al., 2022) | 0.521±0.056 | 0.340±0.060 | 0.741±0.154 |
| | DeepCSR (Cruz et al., 2021) | 0.618±0.103 | 0.397±0.080 | 0.823±0.211 |
| | FreeSurfer (Fischl, 2012) | 0.556±0.049 | 0.364±0.054 | 0.764±0.118 |
| TRT | SurfNet<sub>V</sub> (Ours) | **0.442**±0.016 | **0.218**±0.027 | **0.483**±0.048 |
| | CortexODE (Q. Ma et al., 2022) | 0.457±0.021 | 0.238±0.031 | 0.504±0.071 |
| | DeepCSR (Cruz et al., 2021) | 0.505±0.047 | 0.297±0.053 | 0.610±0.100 |
| | FreeSurfer (Fischl, 2012) | 0.476±0.015 | 0.253±0.022 | 0.519±0.048 |

#### 4.4.2 Cortical Thickness Estimation

In contrast to the alternative methods that rely on the ICP algorithm for registering WM and pial surfaces before calculating the Euclidean distance (Bongratz et al., 2022), our proposed method directly provides vertex-wise cortical thickness estimation. To validate the utility of our estimation, we estimated cortical thickness of 300 subjects (*n*_*AD*_ = *n*_*NC*_ = *n*_*MCI*_ = 100) from the ADNI-2GO (Jack Jr et al., 2008) dataset for comparing our method with FreeSurfer. We computed the average cortical thickness across 35 cortical regions based on the surface parcellation provided by FreeSurfer (Fischl, 2012). Fig. 8 shows the correlation between our thickness estimation and FreeSurfer’s results. In Panel (a), cortical thickness was estimated by calculating the Euclidean distance between corresponding vertices, and its correlation with FreeSurfer’s results was computed. In Panels (b)–(d), cortical thickness was estimated by tracking the deformation trajectory of each vertex. Panel (b) shows the comparison of our estimation with FreeSurfer’s. The significant correlation has demonstrated the effectiveness of our method in capturing cortical thickness, indicating that our method holds promise for quantifying cortical thickness. Panel (c) shows the distribution of correlation values for individual subjects, highlighting consistency in cortical thickness estimation across subjects. Panel (d) further shows the distributions of correlation values across different cortical regions, identifying regions with varying levels of agreement.

**Figure 8:**
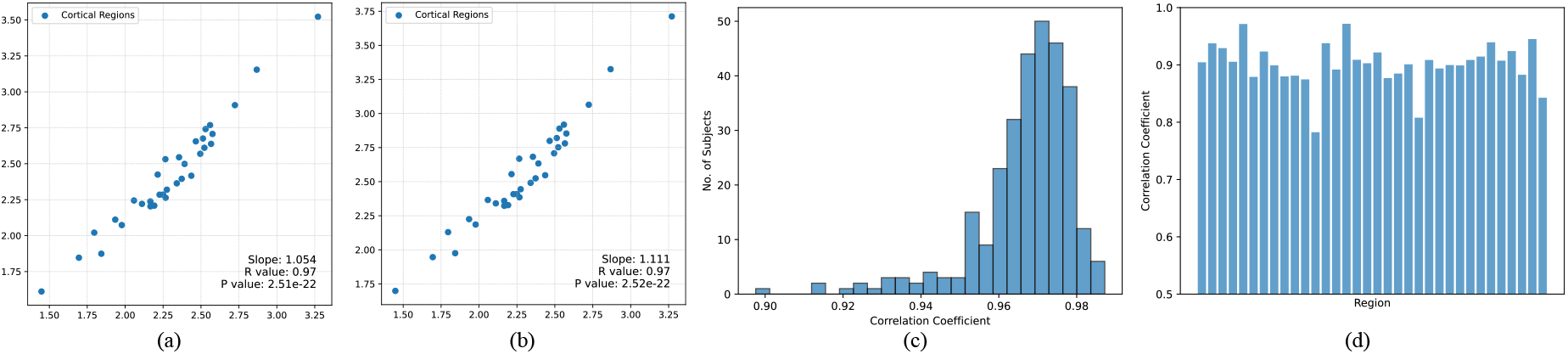
(a) Correlation between cortical thickness estimated by Euclidean distance (Y-axis) and that obtained from FreeSurfer (X-axis) across 35 cortical regions. (b) Correlation between cortical thickness estimated by our network (Y-axis) and that obtained from FreeSurfer (X-axis) across 35 cortical regions. (c) Distribution of correlation values for individual subjects. (d) Distributions of correlation values across subjects within each cortical region.

## 5 Limitations and Future Directions

Despite achieving improved CSR accuracy and a low SIF ratio, our method can be further improved by adopting better segmentation maps for initialization, refining pre- and post-processing methods, and developing new loss functions to minimize the SIF ratio and improve surface quality. While focusing on cortical thickness estimation in this paper, we recognize the value of incorporating other cortical attributes like surface area and sulci depth for characterizing the cortical surfaces, which could serve as complementary constraints. It should be noted that further analysis is needed to thoroughly evaluate the proposed method on diverse cohorts of subjects (e.g., children) although we have demonstrated the correlation between the estimated cortical thickness and that of FreeSurfer on a dataset of Alzheimer’s patients and cognitively normal seniors.

## 6 Conclusion

We have developed a new DL framework, SurfNet, for coupled cortical surface reconstruction and simultaneous cortical thickness estimation. Specifically, the midthickness surface with spherical topology is initialized from the cortical ribbon segmentation maps of each MRI, and optimized to lie at the center of the inner and outer cortical surfaces and deformed to the inner and outer cortical surfaces by three diffeomorphic flows. Two instances of SurfNet based on dense velocity fields and neural ODE are proposed, along with two topology-preserving and inverse-consistent transformation regularizations to enhance diffeomorphism. Extensive experiments on three large-scale datasets have demonstrated the superior performance of our method. Moreover, our method simultaneously estimates cortical thickness, facilitating brain morphometry analyses.

## Footnotes

1 This work has been submitted to the IEEE for possible publication. Copyright may be transferred without notice, after which this version may no longer be accessible.

